# An oomycete effector impairs autophagy in evolutionary distant organisms and favors host infection

**DOI:** 10.1101/697136

**Authors:** Serena Testi, Marie-Line Kuhn, Valérie Allasia, Pascaline Auroy, Fantao Kong, Gilles Peltier, Sophie Pagnotta, Julie Cazareth, Harald Keller, Franck Panabières

## Abstract

An arsenal of effector proteins from plant pathogenic *Phytophthora* species manipulates their host from inside the cells. *Phytophthora parasitica* produces the effector AVH195 during an initial, biotrophic phase of infection. The protein transiently impairs plant immune-associated hypersensitive cell death in *Nicotiana*. ATG8 Interaction Motifs in the protein indicate that the effector targets the autophagic core machinery. We selected a photosynthetic microalga with a single copy *ATG8* gene as an alternative model to dissect AVH195-induced autophagic perturbation. AVH195 slows down autophagic flux in *Chlamydomonas reinhardtii* thus promoting the accumulation of cargo-rich vesicles. In yeast, membrane-associated AVH195 interacts with ATG8 from Chlamydomonas and with different ATG8 isoforms from *Arabidopsis thaliana*. The overexpression of *Avh195* in Arabidopsis promotes growth of both infecting *P. parasitica* and *Hyaloperonospora arabidopsidis*, an obligate biotroph. To our knowledge, this report provides first evidence that an oomycete effector non-selectively targets ATG8 in different organisms from the green lineage to slow down autophagic flux for infection.

## Introduction

Oomycetes have modeled agriculture since the 19th century, and plant protection strategies have still to cope with emerging and re-emerging pathogens belonging to this deep assemblage of filamentous eukaryotic microbes (Kamoun et al., 2015). Oomycetes share with other plant pathogens the ability to possess hundreds of virulence genes encoding effectors that are delivered into plant cells to modulate host cellular processes, defeat defense mechanisms and ultimately promote infection (Gibriel et al., 2016). Effectors from the RxLR class constitute by far the most prominent group in oomycetes. They are generally small (10-25 kDa) proteins without any obvious functional domain, although both predictive and experimental approaches suggest that many of them would adopt common structures, despite substantial sequence diversity (Franceschetti et al., 2017). Deciphering precisely their mode of action requires combined approaches, including expression profiling, comparative analysis of sequences and subsequent three-dimensional structures, as well as the characterization of their molecular targets in the host. The development of combined approaches including-omics and protein-protein interaction tools led uncovering the nature of some plant functions that are targeted by effectors, better understanding the contribution of these functions to plant immunity, and identifying the strategies developed by the oomycete to divert them. Effectors have been shown to perturb essential functions that assure cellular homeostasis and defense mechanisms in the host (Le Febvre et al., 2015). The effector PexRD54 from the potato late blight pathogen, *Phytophthora infestans*, was recently shown to interact with a member of the autophagy-related (ATG) protein family, ATG8. The effector interferes with complex formation between ATG8, which is required for the formation of autophagosome membranes, and Joka2, an autophagic cargo receptor. The interference seems to stimulate pathogen-driven autophagosome formation, possibly to favor elimination of plant defense-related compounds and/or to supply the pathogen with degradation products (Dagdas et al., 2016).

This report is in line with the current knowledge on the emerging role of autophagy in plant-microbe interactions. Autophagy is an universal recycling mechanism that ensures cellular homeostasis through mediating the availability of metabolic building blocks and the repair and turnover of damaged cellular components (Klionsky et al., 2016). In plants, autophagy regulates a range of processes including nutrient management (Masclaux-Daubresse et al., 2017), responses to nitrogen and carbon starvation, and, more generally, stress responses (Avin-Wittenberg et al., 2018). The various functions of autophagy are orchestrated by multifunctional ATG proteins, which are highly conserved among eukaryotes (Meijer et al., 2007). Autophagy also contributes to the recognition of pathogens, and to plant immunity (Hofius et al., 2017). ATG6 (Beclin), a key component of the autophagy machinery, contributes to the restriction of Hypersensitive Response (HR)-associated programmed cell death (PCD) at pathogen infection sites, an effect directly involving the autophagy pathway (Patel and Dinesh-Kumar, 2008). However, this pro-survival contribution of autophagy to plant resistance was further challenged with the characterization of pro-death functions of autophagy during the HR (Hofius et al., 2009). Such apparently contradictory results may be illustrated by the finding that mutants, which are impaired in various ATG genes, display enhanced susceptibility towards necrotrophic pathogens, but exhibit marked resistance against biotrophic microbes (Lenz et al., 2011). Due to the importance of autophagy for plant immunity, it thus seems likely that pathogens evolved mechanisms to target the process in host cells to promote infection, which they may either inhibit or stimulate depending on the parasitic lifestyle (Leary et al., 2018).

*Phytophthora parasitica* Dastur (syn *P. nicotianae* Breda de Haan) is a soilborne oomycete pathogen that has been reported on 255 plant genera in 90 families (Panabières et al., 2016). Like most *Phytophthora* species, *P. parasitica* has a hemi-biotrophic lifestyle, meaning that the microbe invades host tissues initially as a biotrophic pathogen, before it switches to necrotrophy and kills the host. Strains from this species possess a wide array of effectors, including 160-300 RxLR candidates, among which 93 appear conserved in a range of isolates from various origins (Dalio et al., 2018). Here, we describe *Avh195*, an RxLR effector expressed by *P. parasitica* during the biotrophic stage of infection of the model plant Arabidopsis. This effector transiently slows down elicitor-induced, HR-associated cell death. In Arabidopsis plants overexpressing *Avh195*, *P. parasitica* biomass develops significantly stronger than in wild-type plants. In addition, plants are also more susceptible to the biotrophic oomycete *Hyaloperonospora arabidopsidis*. By using heterologous model systems including plants and a photosynthetic microalgae we show that AVH195 interacts with ATG8 and inhibits autophagic flux. We will discuss how these activities of *Avh195* may contribute to pathogenicity of *P. parasitica*.

## Results

### *Avh195* is expressed during the biotrophic phase of *P. parasitica* infection

*Avh195* was characterized as an RxLR effector-encoding sequence in an EST library generated from *P. parasitica* infected plant tissues (Le Berre et al., 2008). The entire transcript encodes a 195-amino acid secreted protein that possesses a 20-aa signal peptide, the canonical RxLR-EER motif, and a 125-amino acid effector domain. Blast searches did not reveal any ortholog, except a 202-aa sequence presenting 66.7% identity with AVH195 in the potato late blight pathogen, *P. infestans* (Fig 1). *Avh195* was not found in EST databases generated from non-infection structures and we thus hypothesized that the RxLR effector contributes to the infection process. To validate this hypothesis, we investigated *Avh195* expression in roots of Arabidopsis plantlets at 3, 10, 30 and 48 hours post inoculation (hpi) with *P. parasitica* zoospores. This time course was previously shown to describe the biotrophic stage of the infection cycle and includes the invasive growth step that corresponds to the switch from biotrophy to necrotrophy at 30 hpi (Attard et al., 2010). *Avh195* mRNA was not detectable in non-invasive stages of the *P. parasitica* life cycle, like motile zoospores, germinated cysts or mycelial cultures (not shown). By contrast, *Avh195* transcripts accumulated in the early steps of infection, then slowly decreased during later steps, and became barely detectable at 48 hpi (Fig 2). To further dissect the infection process, we monitored expression of the genes encoding Haustorium-Specific Membrane Protein 1 (HMP1) and Necrosis-inducing *Phytophthora* Protein 1 (NPP1) in *P. parasitica*, which are considered as markers for biotrophic and necrotrophic stages of *Phytophthora* infection, respectively (Jupe et al., 2013). Based on their respective mRNA levels (Fig 2), we concluded that *Avh195* transcripts preferentially accumulate during the biotrophic stage of the *P. parasitica* infection cycle.

**Figure 1:**
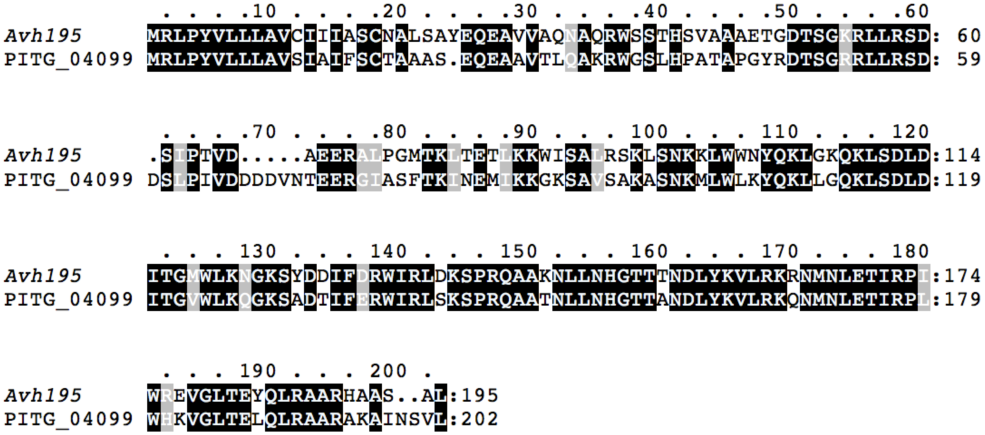
Alignment of AVH195 and PITG_04099. The amino acid sequences were aligned using Clustal Omega and edited using Matchboxshade for Macintosh. Shading indicates blocks of identical (black) or similar (grey) amino acids.

**Figure 2.**
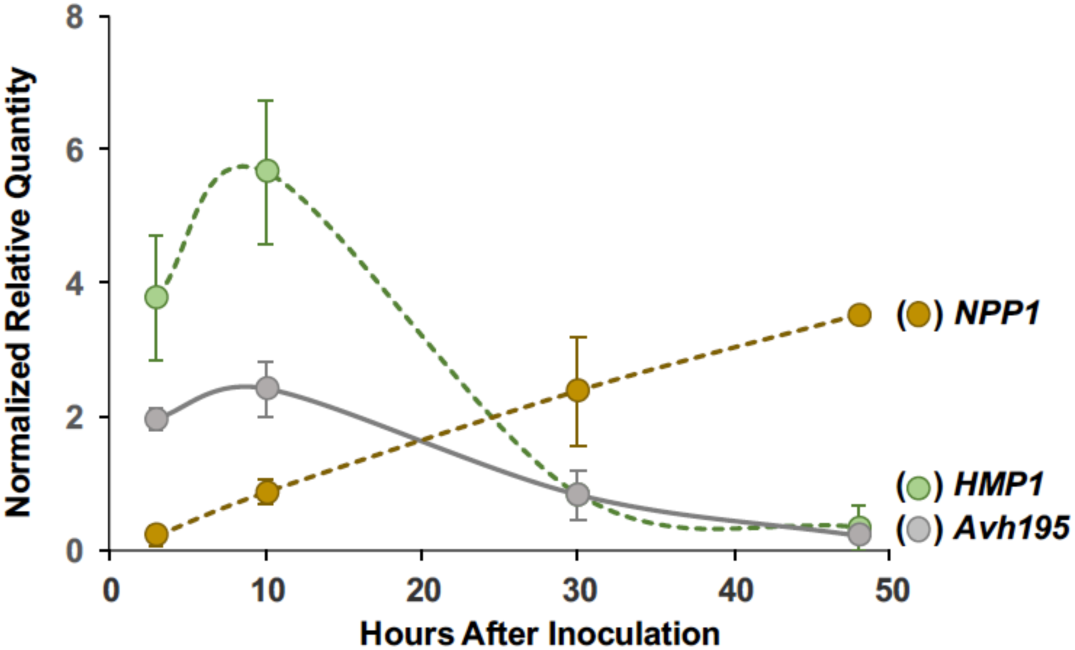
Expression profile of Avh195 during the P. parasitica infection cycle. Relative *Avh195* mRNA levels (grey) were determined by quantitative RT-PCR in *A. thaliana* plantlets inoculated with *P. parasitica* zoospores at different time-points of infection. The expression profiles of *HMP1* (green) and *NPP1* (brown) describe the biotrophic and necrotrophic stages of the infection, respectively. Relative transcript quantities were normalized with transcripts from the *UBC* and *WS41* reference genes. Presented are the means (± SD) from 3 biological replicates.

### *Avh195* has the potential to interfere with autophagy

Examination of the AVH195 protein sequence revealed 5 potential ATG8 interacting motifs (AIMs), otherwise designed as LC3-Interacting regions, or LIRs (Jacomin et al., 2016). AIMs consist of 6 amino acids that are required for the interaction with ATG8 (Birgisdottir et al., 2013). AIMs were detected in the AVH195 sequence by comparison with a collection of LIR motif-containing proteins (LIRCPs) from various organisms that are compiled at the iLIR autophagy database. The AIMs of LIRCP are defined by a position-specific scoring matrix (PSSM), which is derived from the alignment of experimentally verified AIM/LIR motifs (Kalvari et al., 2014). The scores for AVH195 ranged from 7 to 16 (Figure 3). By setting the score cut-off to 10, we retained 3 candidate AIMs for further analyses.

**Figure 3:**
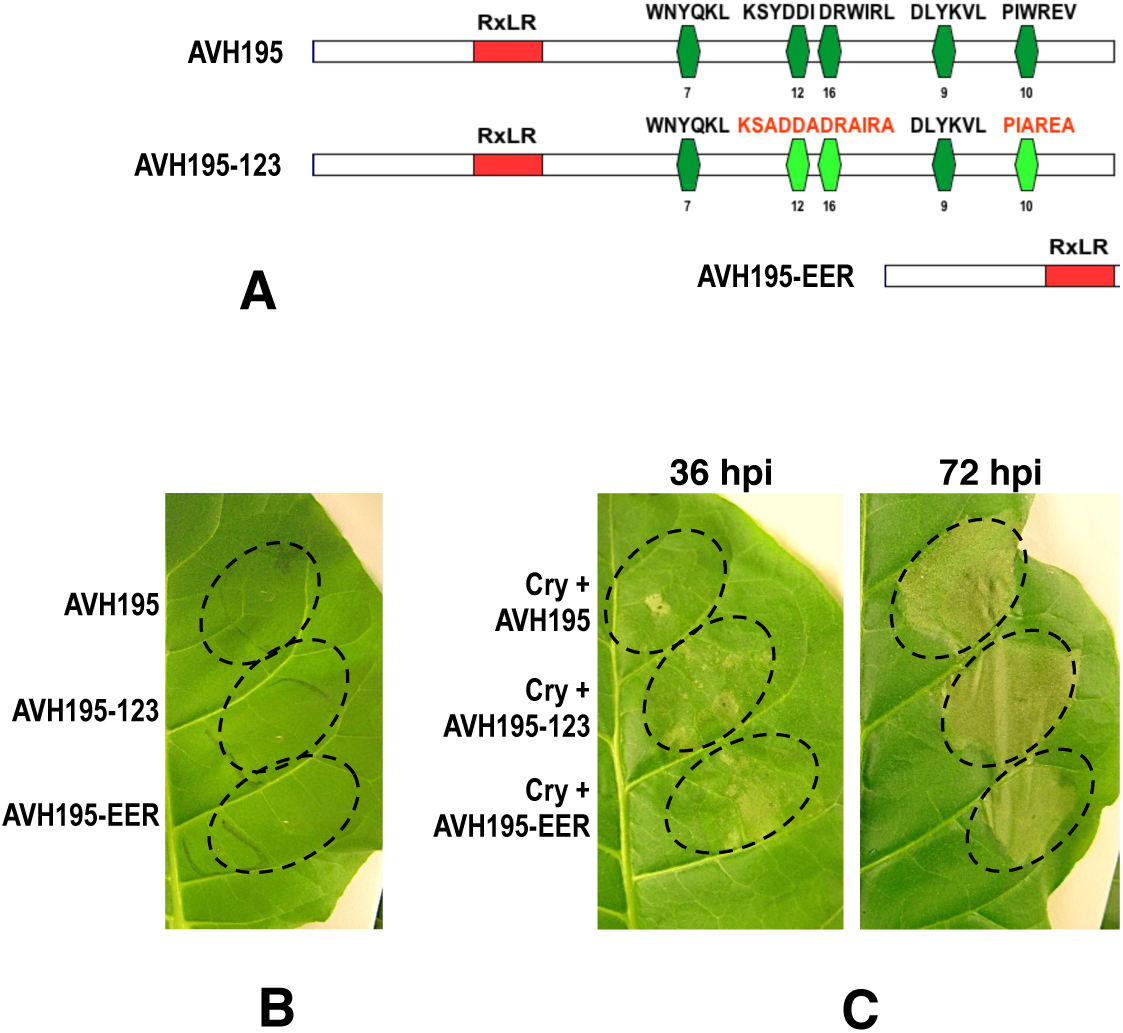
AVH195 requires AIMs for transient inhibition of the HR. **A.** Graphic representation of the different constructs used for agroinfiltration experiments. AVH195: The entire ORF of *Avh195* without the signal peptide. AVH195-123: Three AIMs highlighted in bright green were deleted by site-directed mutagenesis. AVH195-EER: The negative control, in which Avh195 is devoid of the entire effector domain. Numbers below the indicated AIMs represent the Position-Specific Scoring Matrix values. **B.** Agroinfiltrations of *N. tabacum* leaves with the three constructs provoke no visible symptoms at 36h post inoculation. **C.** Tobacco leaves were co-transfected with *A. tumefaciens* strains harboring the cryptogein gene and each of the three gene constructs. Symptoms were observed at 36 hpi (left) and 72 hpi (right). Shown is a representative experiment from three independent replicates, each involving 5 plants and 4 infiltrated leaves per plant.

### *Avh195* transiently antagonizes HR-associated cell death in tobacco

We assessed the possible contribution of *Avh195* to *P. parasitica* virulence through transient expression in *N. tabacum* leaves via *A. tumefaciens* carrying various forms of the effector (Fig 3). They consisted in the entire ORF of *Avh195* (without the signal peptide), a construct in which hydrophobic residues of the three retained AIM sites were changed into alanine (195-123) and a construct deleted of the entire C-terminal effector domain beyond the RxLR-EER motifs (195-EER, Fig 3A). No cell death was observed in any case, indicating that AVH195 has no toxic effect and/or does not trigger an HR in tobacco (Fig 3B). We then checked the ability of the effector and its variant forms to interfere with plant cell death by co-infiltration with a construct expressing cryptogein, an elicitin secreted by *P. cryptogea* that induces strong HR-associated cell death in tobacco leaves (Ponchet et al., 1999). Cell death symptoms were significantly reduced, but not abolished, at 36 hpi when cryptogein was co-produced with the intact form of AVH195. This was not the case when cryptogein was co-produced with the mutant variants of AVH195, 195-123 and 195-EER (Fig 3C, left). Yet, further observations revealed that differences in phenotypes started to merge from 48hpi on, and that co-infiltration with all 3 constructs resulted in comparable extents of cell death lesions at 72hpi (Fig 3C, right). Together, these findings show that AVH195 transiently reduces HR cell death. As this interference with cell death relies on functional AIMs, we suggest that a physical interaction between AVH195 and ATG8 contributes to the effect.

### Selection of an alternative model for the functional characterization of *Avh195*

*ATG8* occurs as a multigene family in plants (Kellner et al., 2017). Arabidopsis and tomato encode nine and seven ATG8 members, respectively (Seo et al., 2016). Arabidopsis sequences are further clustered into three subfamilies or clades, display differential expression in various tissues and differentially respond to environmental cues (Slavikova et al., 2008). We thus anticipated that analyses of *ATG8* expression would be difficult to conduct in Arabidopsis. We then decided to assess the perturbation of plant autophagy by *Avh195* on a reduced-complexity model and chose the unicellular photosynthetic alga *Chlamydomonas reinhardtii* for several reasons. First, it is a model organism for studies on a large array of fundamental cellular processes, including development, reproduction, chloroplast biology and photosynthesis (Harris, 2001). Second, techniques and molecular tools are available for this organism, otherwise designed as the “photosynthetic yeast” (Rochaix, 1995). Third, autophagy can be easily induced in Chlamydomonas using a range of conditions, among which treatment with the macrolide rapamycin (Crespo et al., 2005). A fourth and main advantage for experimental studies is that the Chlamydomonas ATG core machinery is encoded by single-copy genes (Avin-Wittenberg et al., 2012).

### AVH195 physically interacts with ATG8

Before investigating a possible interaction between AVH195 and ATG8, we analyzed whether both proteins localize to the same subcellular compartments in plant cells. We generated gene constructs that allowed co-expression experiments in *N. benthamiana* between N-terminal green fluorescent protein (GFP)-tagged AVH195 without the signal peptide and C-terminal red fluorescent protein (RFP)-tagged ATG8 from *C. reinhardtii* (CrATG8). Confocal imaging showed that the GFP-AVH195 fluorescence was restricted to the borders of fully turgescent epidermis cells, while ATG8-associated RFP fluorescence was distributed to the cell border, but also to cytoplasmic filaments and the nucleus (Fig 4A, upper row). The localization pattern of both proteins became more evident in plasmolyzed epidermal cells (Fig 4A, bottom row). Here, AVH195 labeled the tonoplast and vesicle-rich, membrane-surrounded cytoplasmic filaments. RFP-ATG8 located to the cytoplasm and the nucleus and, to a lower extent than AVH195, associated with vesicle-rich structures (Fig 4A, bottom). Merged micrographs revealed several cases of co-localization of both proteins (Fig 4A, right panels).

**Figure 4:**
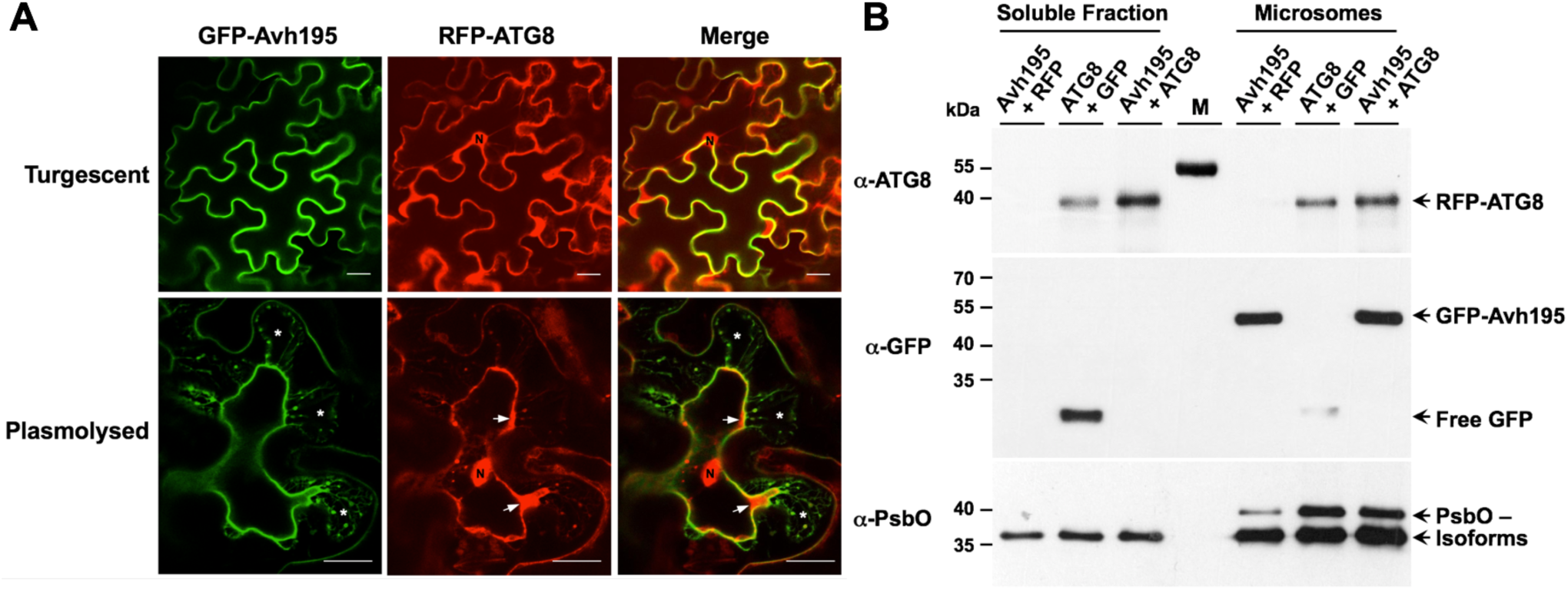
AVH195 localizes to membranes, while ATG8 is both soluble and membrane-associated. **A.** Optical sections of *N. benthamiana* epidermal cells transiently co-expressing *Avh195* and ATG8 from *C. reinhardtii*, as analyzed by confocal laser-scanning microscopy. Analyzed cells were either fully turgescent (upper lane) or plasmolysed (bottom lane) to detect the N-terminal GFP-tagged AVH195 protein (left panel), the N-terminal RFP-tagged ATG8 (middle panel), or co-localization of both as yellow color in merged channel micrographs (right panel). Arrowheads point to cytoplasmic localization of ATG8. Asterisks in the left and right panels indicate vesicle-rich, membrane-surrounded cytoplasm filaments that attach to the cell wall. N = nucleus. Bars represent 20 µm. **B.** Immunoblotting of soluble and microsomal fractions prepared from *N. bentamiana* leaves transiently co-expressing GFP-tagged AVH195 with free RFP, RFP-tagged ATG8 with free GFP, and GFP-tagged AVH195 with RFP-tagged ATG8. Soluble and membrane-associated proteins were revealed with antisera directed against ATG8 and GFP. An antibody recognizing the photosystem II PsbO protein was used for loading control. Note that the ATG8 antiserum recognizes the 55 kDa prestained protein of the PageRuler Protein Ladder (M).

To support co-localization of AVH195 and ATG8, soluble and microsomal proteins were prepared from tobacco leaves expressing both GFP-AVH195 and RFP-CrATG8, and were analyzed by Western blotting. An anti-ATG8 antiserum detected RFP-CrATG8 in almost equal amounts in both soluble and microsomal fractions (Fig 4B). By contrast, GFP-tagged AVH195 was solely detected in the membrane-associated protein fraction. These findings confirm the microscopic observation that a portion of CrATG8 co-localizes with AVH195 at membranes.

To investigate a potential membrane-associated interaction between AVH195 and ATG8, we used the mating-based split-ubiquitin yeast two-hybrid system (mbSUS), which has been developed to detect the interactions between membrane-anchored proteins and their partners (Grefen et al., 2009). Yeast transformants expressing the native *Avh195* bait with the CrATG8 prey readily grew on selective medium, indicating physical interaction between the two proteins (Fig 5A). We then tested three members of the *ATG8* family from Arabidopsis. We selected *AtATG8D*, *AtATG8G* and *AtATG8H* as representatives of the three established clades (Seo et al., 2016). Yeast cells expressing the native AVH195 bait and the AtATG8 preys were able to develop on selective medium, although vigor of growth was strongest with AtATG8H as the prey and weaker with AtATG8D and AtATG8G (Fig 5A). Yeast cells expressing the triple AVH195 AIM mutant as bait were severely compromised for growth on selective medium (Figure 5B). Taken together, these results indicate that AVH195 associates with ATG8 in membranous compartments of the cell, and that this interaction is mediated by the canonical AIMs of the effector.

**Figure 5:**
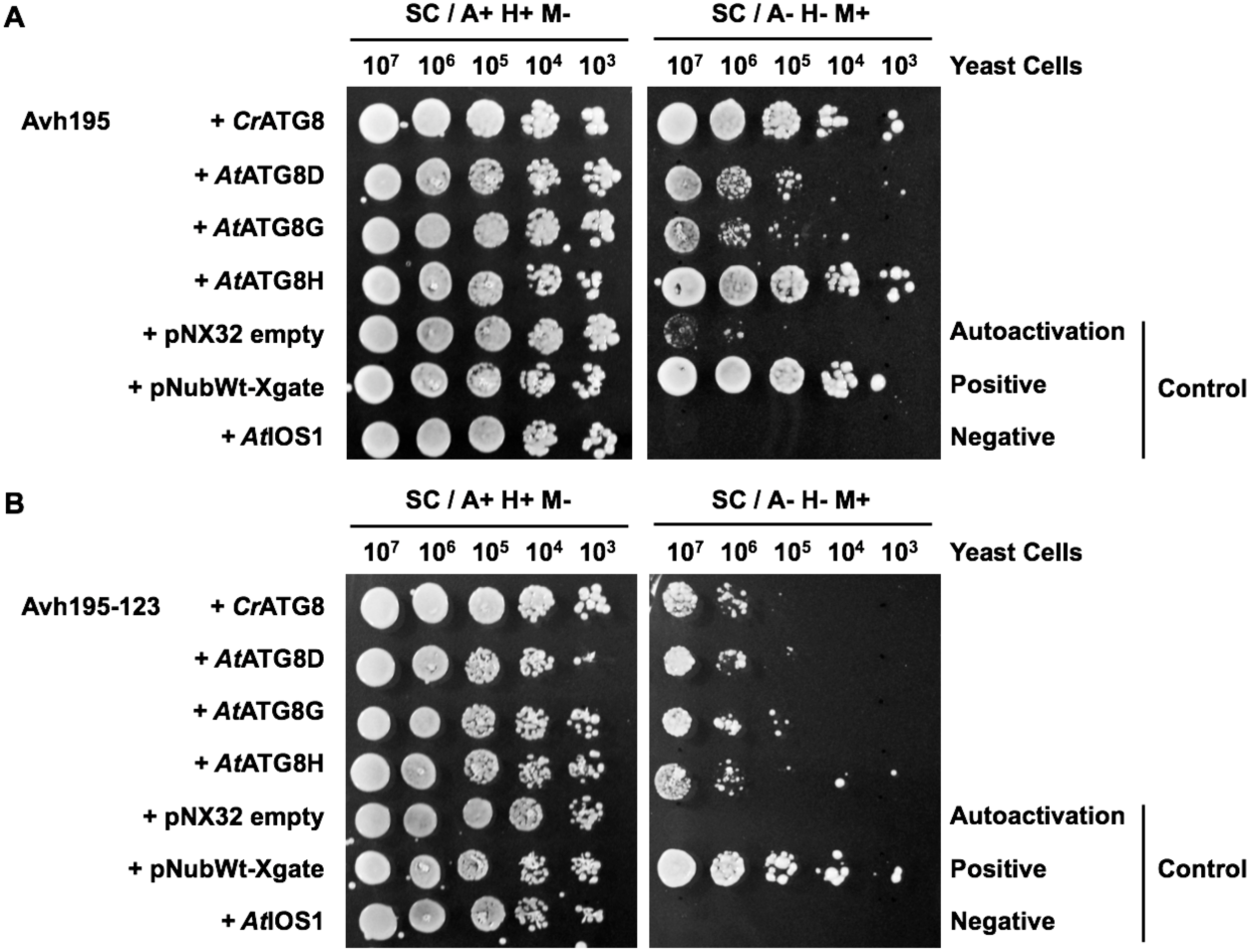
AVH195 requires AIMs for interaction with ATG8. The mating-based split-ubiquitin yeast two-hybrid system was used to monitor the interaction of membrane-anchored AVH195 with ATG8 from *C. reinhardtii* and *A. thaliana*. **A.** Coexpression of the native AVH195 bait with ATG8 preys allows yeast cells to grow on selective medium in the absence of adenine (A) and histidine (H), and in the presence of methionine (M). **B.** The mutation of 3 AIMs in the AVH195 bait variant (AVH195-123) decreases the capacity of yeast cells to grow on selective medium. The negative prey control *AtIOS1* encodes an Arabidopsis receptor.

### Heterologous expression of *Avh195* in Chlamydomonas does not impact vital fitness parameters

To estimate whether AVH195 interferes with the core autophagic process, we transformed *Chlamydomonas reinhardtii*. The *C. reinhardtii* nuclear genome has a GC content of ∼ 65% (Merchant et al., 2007). We thus designed a synthetic *Avh195* coding region (without the signal peptide) with a codon usage adjusted to the algae, that was introduced into the cell line *dw15.1*, which is devoid of a cell wall (see Methods). Transformants were first selected for the presence of the transgene on a set of 101 independent lines. A further selection step assessed the expression of the transgene using RT-qPCR on a total of 8 independent transformant lines revealed that the relative mRNA level of *Avh195* greatly varied from one transformant to an other, likely due to positional effects (Cerutti et al., 1997). We retained three independent lines, N15, N26, and C40 for all further analysis.

To characterize a potential impact of transgene expression on the global fitness of the transformants, we synchronized the cultures by alternating light and dark periods over 24-hours. We stimulated autophagy by supplementing cultures with 0.5 µM rapamycin, a concentration that does not affect vital parameters of Chlamydomonas (Crespo et al., 2005). We avoided using higher concentrations of rapamycin, which induce drastic changes in synchronized cultures, such as a marked decrease in cell number, an increase in the mean cell volume, and an arrest of proliferation (Juppner et al., 2018), to analyze subtle phenotype changes that we expected to be induced by *Avh195*.

To assess effects of transformation on Chlamydomonas physiology, we analyzed several vital parameters by spectral flow cytometry over a circadian cycle of 24h. Cell size was measured by forward light scattering (FSC), and the complexity of cellular structure by side light scattering (SSC). Cell proliferation rate was assessed by measuring the dilution of the fluorescence intensity of 5-,6-carboxyfluorescein succinimidyl ester (CFSE) upon cell division (Lyons, 2000). The addition of 4′,6-diamidino-2-phenylindole (DAPI) to the cells allowed evaluating cell death. Finally, autofluorescence of the cells reflected potential changes in the chlorophyll content of cultures. As a rule, the different parameters were tightly connected to the circadian cycle. Cell proliferation predominantly occurred during the night, but stagnated under light, whereas all other parameters (including DAPI) reached a peak after 12h of light and decreased during the night until reaching a level similar to that observed at the beginning of the analysis (T_0_, Supplementary Figure S1). Wild-type (Wt) and transformant cultures increased cell number with similar kinetics and to similar magnitudes over the light period and decreased likewise. In addition, no significant difference was observed between untreated and rapamycin-treated cultures (Supplementary Figure S2), confirming that the mild doses of the drug do not affect vital parameters of Chlamydomonas cells (Crespo et al., 2005). Only slight differences between cell lines were observed, which likely reflected individual variations rather than a global trend. We thus concluded that transformation and expression of *Avh195* had no incisive effects on the physiology of the cultures.

### AVH195 slows down autophagy in *Chlamydomonas reinhardtii*

We then compared the subcellular organization of Wt and transformed Chlamydomonas cultures using transmission electron microscopy (TEM). The major difference between untreated cells that express or do not express *Avh195* was the accumulation of starch granules to obvious higher amounts in transgenic lines (Fig 6A, left column). Starch content in Chlamydomonas is under tight control of the circadian clock, reaching a minimum in the middle of the light phase at 6h in a 12h diurnal rhythm (Ral et al., 2006). We thus compared the average number of starch granules per cell in Wt and transgenic cultures that were supplemented or not with 0.5 µM rapamycin. The measurements were performed at 4h and 8h after the beginning of the light phase (and after addition of rapamycin), to evaluate starch accumulation at the end of the degradation phase and the beginning of synthesis. The number of granules was significantly higher in cells from the three transformants, when compared to Wt cells, both at the end of degradation and the beginning of synthesis (Fig 6B). Increased starch accumulation in *Avh195*-expressing cells was independent of rapamycin treatment, as no significant difference was observed between untreated and treated cultures (Fig 6B).

**Figure 6:**
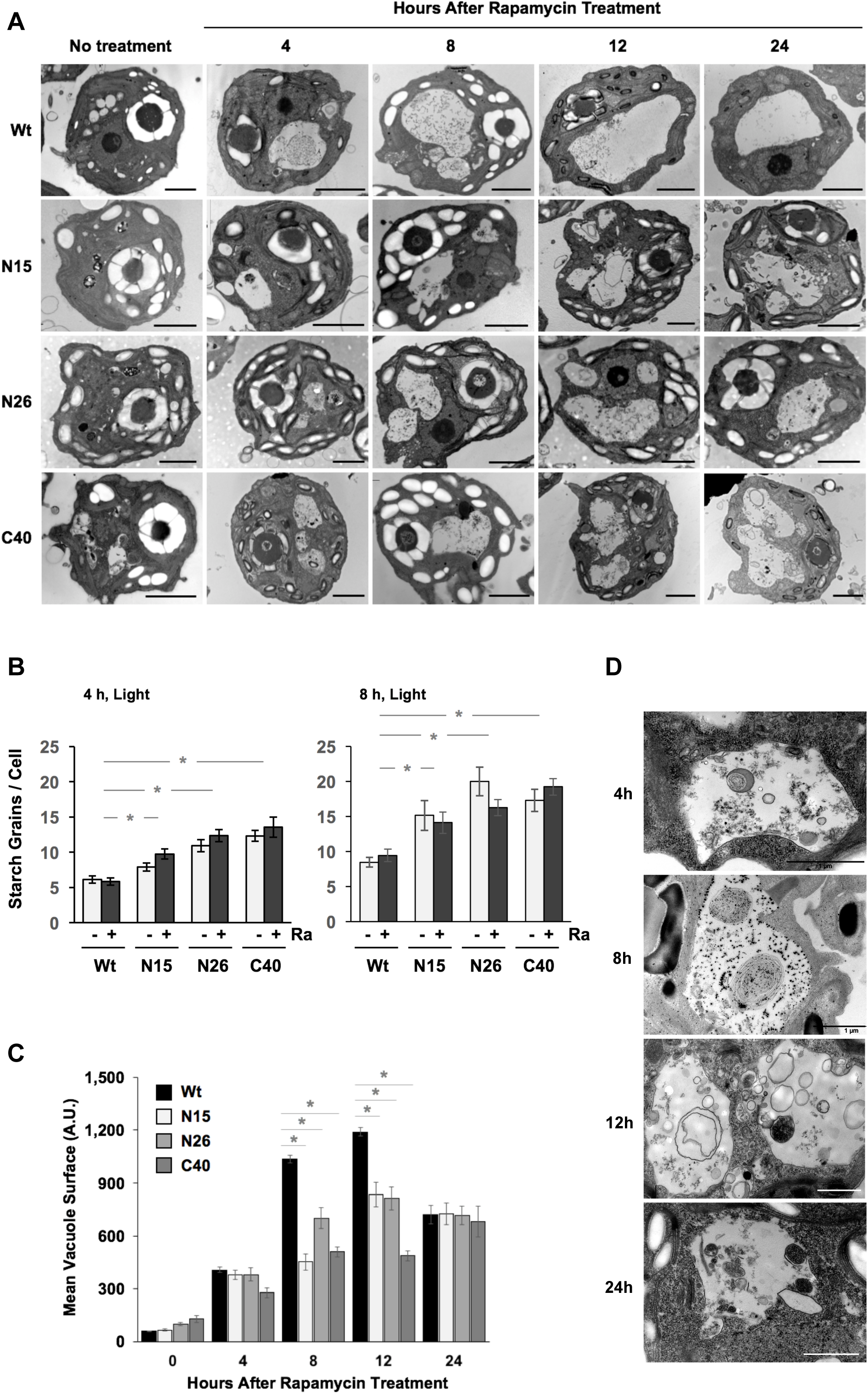
Subcellular phenotypes of *Chlamydomonas* cells expressing *Avh195*, as analyzed by transmission electron microscopy. **A.** Under non-induced conditions, wild type (Wt) cells (upper lane) contain several empty vacuoles that fuse upon the addition of rapamycin to form a predominant, large vacuole 4 h after onset of treatment. Cells overexpressing *Avh195* display an accumulation of nondigested material in lysosome-like structures, starch accumulation, and a delay in central vacuole fusion, when compared to the Wt. No obvious other modifications in the subcelluar organisation are related to effector expression. The bars represent 2 µm. **B.** *Chlamydomonas* transformants expressing *Avh195* accumulate starch grains to significantly higher amounts than the Wt, independent of rapamycin treatment. The number of starch grains was determined 4 h and 8 h after onset of the light period and rapamycin (Ra) treatment on micrographs from 5 independent TEM sections representing each 10-12 cells per treatment and time point. Shown are means (±SE) from n=25 cells per line and treatment. **C.** Rapamycin-induced vacuole swelling is impaired in *C. reinhardtii* transformants expressing *Avh195*. In the *Avh195*-expressing lines, the predominant vacuole is replaced by smaller vacuoles. For each line and time point of treatment, micrographs from 4-5 independent TEM sections representing 10-12 cells each were analyzed for the presence of vacuoles. The surface of the biggest vacuole in each cell was then determined using the ImageJ software. Shown are means (±SE) from n=30 vacuoles per line and time point of treatment. A.U. = Arbitrary Units. Asterisks in **B** and **C** indicate significant differences with p<lt;0.05, as determined by Student’s t-test. **D.** Higher resolution views of *Avh195*-expressing cells from line N26 presenting lysosome-like vacuoles containing electron dense material. This material started to accumulate in vacuoles 4 h after rapapycin treatment. The undegraded cargo persists without elimination over 24 h of autophagy induction. Bars represent 1 µm.

Another rapamycin-independent difference between Wt and transformant cultures was observed at higher TEM resolution. *Avh195*-expressing cells accumulated electron-dense, lysosome-like structures, which did not evolve over the 24h-day/night cycle (Supplementary Figure S3). This observation suggested that AVH195 interferes with basal autophagic flux in the transformant cells. Upon rapamycin treatment, empty vacuoles fused to form a predominant, large vacuole in Wt cells as soon as 4 h after onset of treatment (Fig 6A, upper lane and Supplementary Figure S4). Conversely, cells overexpressing *Avh195* displayed a marked accumulation of nondigested material in lysosome-like structures, and a delay in central vacuole fusion, when compared to the Wt. At 24h after rapamycin treatment, this delay in central vacuole formation was no longer obvious (Fig 6A, right column and Supplementary Figure S4). To determine the dynamics of vesicle fusion and the formation of a central vacuole in a quantitative manner, we measured the surface of the biggest vacuole in individual cells on 4 to 5 TEM micrographs that represent about 10 cells each. The surface of vacuoles in Wt cells increased constantly to maximum sizes at 12h after rapamycin treatment (Fig 6C). By contrast, cells from the 3 transgenic lines appeared impaired in vacuole merging, as they maintained significantly smaller vesicles over the initial 12h of treatment. The differences in vacuole size between the Wt and the transgenics were no longer observed after 24h of treatment (Fig 6C). At higher TEM resolution, transformant cells displayed an accumulation of various cytoplasmic debris and organellar-like structures over time of rapamycin treatment (Fig 6D). We assumed that these structures were autophagic bodies and their debris, which accumulate in vesicles of the transformant cells to witness impaired autophagic flux.

Observations similar to what we describe for *Avh195*-expressing cells were previously made in Chlamydomonas cells that were treated with Concanamycin A, a drug that inhibits autophagic flux (Couso et al., 2018). We thus attempted to validate this hypothesis by measuring the accumulation of soluble and membrane-bound ATG8 upon treatment of Chlamydomonas cultures with rapamycin and Concanamycin A. Under control conditions (no supplementation with drug), Wt and transformants cultures displayed a similar pool of soluble (inactive) ATG8, whereas membrane-associated (active) ATG8 was barely detectable in Western experiments (Figure 7, left panel). Rapamycin treatment induced a marked accumulation of two distinct forms of membrane-associated ATG8, which was stronger in cells from the transformant lines than from the Wt (Figure 7, middle panel). By contrast, we did not observe any change in the pool of soluble ATG8, whatever cell line analyzed. Treatment with Concanamycin A also induced the accumulation of two forms of active, membrane-associated ATG8 thus reflecting the impairment of autophagic turnover of the protein (Figure 7, right panel). Taken together, our results from TEM and molecular analyses indicate that AVH195 interacts with ATG8 to delay the autophagic turnover in Chlamydomonas, which is illustrated by delayed formation of vacuoles, the inability of small vesicles to fuse with vacuoles, the accumulation of cellular debris within vesicles, a marked accumulation of starch granules, and by an increased accumulation of membrane-associated ATG8. We therefore suggest that AVH195 delicately alters the dynamics of the autophagic flux, rather than blocking the process completely.

**Figure 7:**
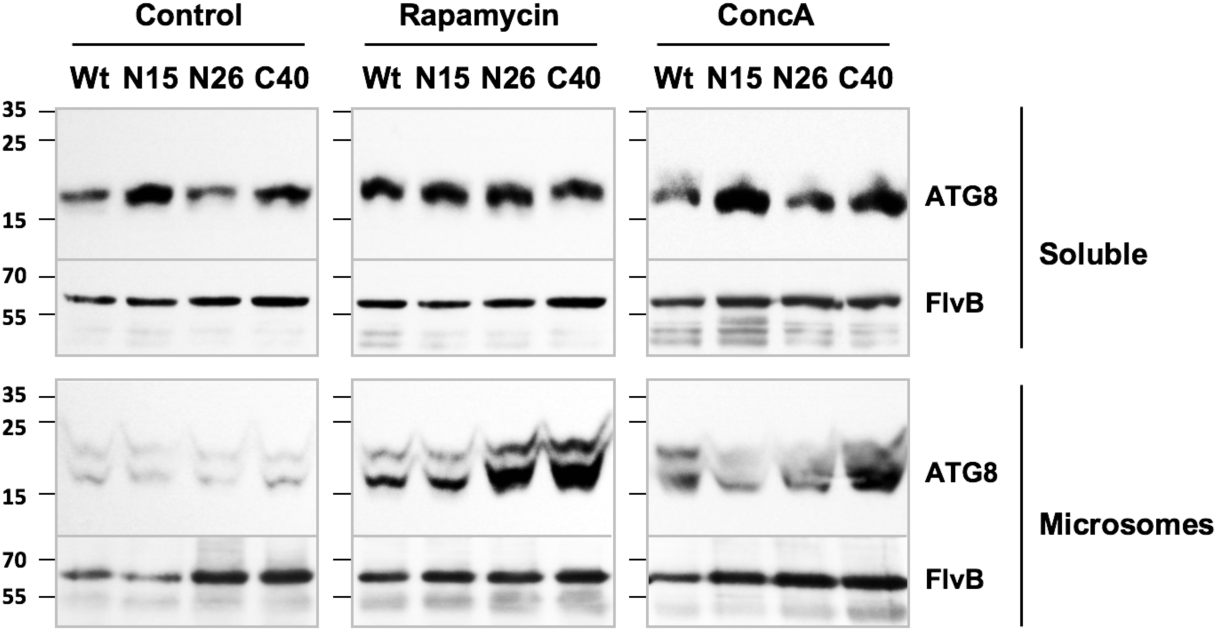
AVH195 delays autophagic turnover of membrane-bound ATG8 in *C. reinhardtii*. Cell suspensions of the wild-type (Wt) and the transgenic lines N15, N26, and C40 were supplemented with solvent alone (Control, left column), and with Rapamycin (middle column) or Concanamycin A (ConcA, right column), and incubated for 8 h under light before being harvested. Protein extracts from the individual cultures were separated into soluble and microsomal fractions, and subjected to SDS-PAGE followed by immunoblotting and detection with the ATG8 antibody. An antibody directed against the chloroplast *C. reinhardtii* flavodiiron B (FlvB) protein was used for normalization.

### Expression of *Avh195* in Arabidopsis promotes infection by oomycete pathogens

We generated transgenic Arabidopsis lines (ecotype Col-0) harboring the *Avh195* coding region under the control of the constitutive Cauliflower Mosaic Virus (CaMV) *35S* promoter. We selected 2 independent transgenic lines that accumulate significant amounts of effector transcripts. Arabidopsis is non-responsive to rapamycin (Mahfouz et al., 2006), but plant autophagy is activated for energy production at night (Izumi et al., 2013). To determine whether the observed effect of Avh195 on autophagic turnover in Chlamydomonas is also observed in plants, we prepared soluble and microsomal protein fractions from Wt and *Avh195*-expressing Arabidopsis plants that were kept in the dark. Under these conditions, Wt Arabidopsis accumulated only marginal amounts of membrane-associated ATG8, showing that autophagic flux is functional in these plants at night (Fig. 8A, right panel). By contrast, plants from the transgenic lines accumulated larger quantities of membrane-associated ATG8, indicating that autophagic turnover is hindered in plants expressing *Avh195* (Figure 8A, right panel).

**Figure 8:**
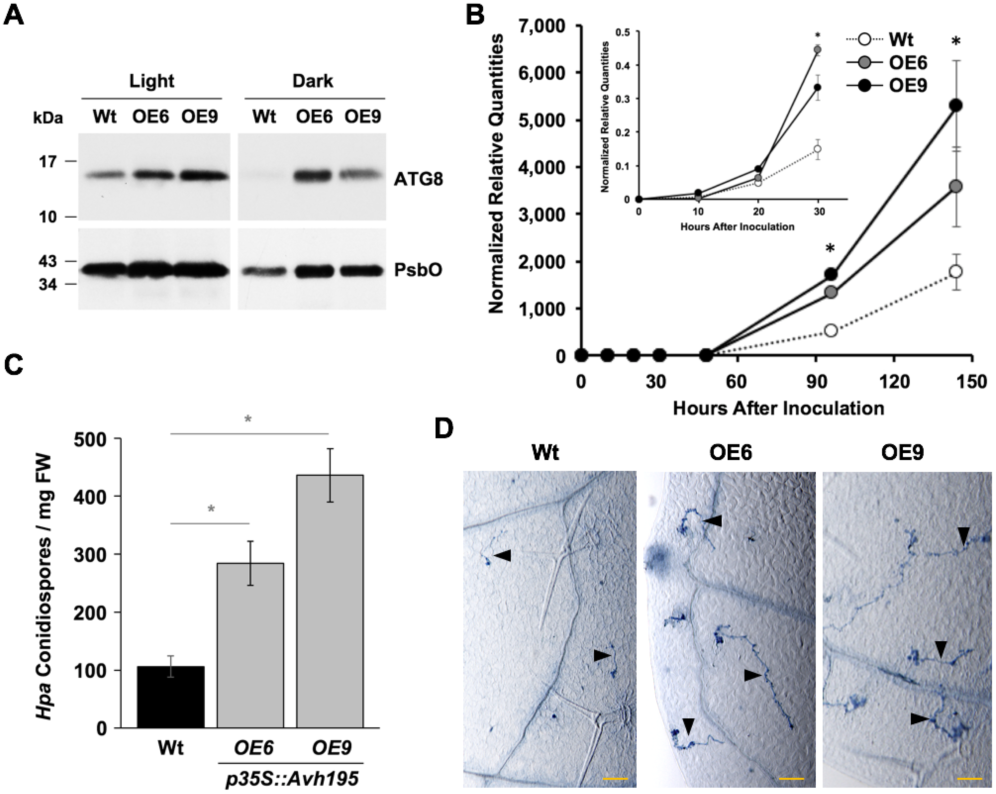
Expression of *Avh195* in transgenic *A. thaliana* alleviates ATG8 turnover and increases susceptibility to Oomycete pathogens. **A.** Accumulation of ATG8 in microsomal fractions from the wild-type (Wt) and from 2 independent transgenic lines expressing *Avh195*. In the dark, transgenic lines maintain higher quantities of membrane-associated ATG8 than the Wt. An antibody recognizing the PsbO protein was used for loading control. Note that the PsbO antibody reveals a single band in microsomal extracts from Arabidopsis, whereas two distinct isoforms are detected in *N. benthamiana* (Figure 4). **B.** Biomass of *P. parasitica* develops faster and stronger in *Avh195*-expressing plants, when compared to the Wt. RNA extracted from inoculated samples were analyzed by RT-qPCR for the accumulation of transcripts from the constitutively expressed *P. parasitica* gene *WS41*. The insert shows the evolution of biomass during the first 30 hours post inoculation in the same experimental onset. Shown are the means (±SD) from 3 replicates. Asterisks indicate significant differences with p≤0.05, according to Student’s t-test. **C, D**. *Avh195-*expressing plants are more susceptible to the downy mildew pathogen, *Hyaloperonospora arabidopsidis* (*Hpa*). **C.** Conidiospore production on plants from the *Avh195*-overexpressing lines increases at least 3-fold, when compared to wild-type plants. Bars represent mean values ±SE for 20 samples. Statistically significant differences with p≤0.001 were determined by Student’s t-test and are indicated with an *. The graph shows a representative experiment among 3 repetitions. **D.** Invading *Hpa* hyphae (arrowheads) develop better in *Avh195-*expressing plants, when compared to the Wt. Infected leaves from the Wt and the transgenic lines were collected 24 h post inoculation and stained with trypan-blue. The micrographs show representative focused sections (see Supplementary Figure 5 for overview). Bars represent 100 µM.

To assess the consequences of *Avh195* expression on pathogen development, seeds from homozygous transgenic lines and from the Wt were inoculated with zoospores from *P. parasitica* (see Methods). The development of the oomycete within plant tissues was then determined by RT-qPCR using samples of infected plants that were collected 10h, 20h, 30h, 48h, 96h, and 144h after inoculation. We correlated the development of oomycete biomass within plant tissues with the expression of the *P. parasitica* gene *WS41,* which is widely accepted as being constitutively expressed during plant infection (Attard et al., 2014; Yan and Liou, 2006). *Phytophthora* transcripts became clearly detectable at 20hpi and accumulated to significantly higher levels in the two transgenic lines, when compared to the Wt (Fig 8B). The differences in biomass increase between the transgenic lines and the Wt were still detected at 144 hpi. We thus concluded that plants expressing *Avh195* were significantly more susceptible to *P. parasitica* than Wt plants. To test whether AVH195 confers susceptibility to an oomycete with a different infection strategy, we inoculated plants from the transgenic lines and from the Wt with the obligate biotrophic pathogen, *Hyaloperonospora arabidopsidis* (*Hpa*). To estimate the infection success, we determined the amount of asexual conidiospores that are produced at the end of the infection cycle (Hok et al., 2011). Sporulation rates were significantly higher in plants from the transgenic lines, when compared to the Wt (Figure 8C). To determine whether the increase in sporulation rates on the transgenic lines was correlated with an increased infection success, we analyzed the extent of hyphal growth in leaves at an early stage of infection (24 hpi) in plants from the Wt and the transgenic lines. In both transgenic lines, initial hyphae developed much better than in the Wt (Figure 8D and Supplementary Figure S5). Taken together, our data strongly suggest that AVH195 slows down autophagic flux through interaction with host ATG8 to promote biotrophic growth of oomycete pathogens.

## Discussion

In this study, we show in different photosynthetic systems that the *P. parasitica* AVH195 effector interferes with autophagy through interaction with ATG8. The effector slows down autophagic flux to favor pathogen development within host tissues. To decipher the precise mechanisms underlying the modulation of autophagy by AVH195, we developed functional analyses on the green algae *C. reinhardtii*. To our knowledge, we provide the first report that an oomycete effector is active in a non-plant, but photosynthetic organism, making Chlamydomonas a promising model for further effector studies. The only other trans-kingdom activity has been shown for a CRN effector of the legume root pathogen *Aphanomyces euteiches.* This effector has been successfully expressed in *Xenopus laevis* embryos, where it triggers aberrant developmental modifications (Ramirez-Garces et al., 2016). Developmental phenotypes were not found in the present study, as Chlamydomonas cells expressing *Avh195* do not present modifications other than the perturbation of autophagic flux and the accumulation of starch granules.

Our analyses conducted on both Chlamydomonas and Arabidopsis revealed accumulation of membrane-associated ATG8, as revealed by western blot experiments. Such accumulation might be a consequence of autophagy induction, but also reflects impaired autophagic flux through a defect in vacuolar degradation (Cheong and Klionsky, 2008). Our study on Chlamydomonas cells expressing or not expressing *Avh195* indicated that autophagic flux is transiently impaired. TEM analyses revealed the presence of smaller vesicles in transformants, when compared to Wt cells. Transformants display a delayed progression of autophagy upon rapamycin treatment. They accumulate cellular debris similarly to what is observed in Chlamydomonas cells that are treated with Concanamycin A, an inhibitor of autophagic flux. We also noted increased accumulation of starch in transformants that is illustrated by an increase in starch granules in the cells. Starch has been shown to accumulate in plants treated with 3-methyladenine (3-MA), an autophagy inhibitor, as well as in *atg* mutants (Wang et al., 2013). All our convergent observations strongly suggest that AVH195 negatively modulates autophagy in a transient manner.

Studies of the molecular processes underlying programmed cell death during the HR revealed that autophagy contributes to this mechanism (Liu et al., 2005). A particular role for autophagy was shown in the delimitation of cell death, as *atg6* mutants are unable to restrict spreading of the HR induced by viral and bacterial pathogens (Patel and Dinesh-Kumar, 2008). However, other studies rather revealed a ‘pro-death’ function of autophagy in the execution of HR triggered by avirulent pathogens in Arabidopsis (Hofius et al., 2009). This apparent contradiction was evoked in a number of additional reports, indicating a contribution to the outcome of plant-microbe interactions more subtle than supposed. Slight modulations of this fine-tuned machinery might thus have consequences for success or failure of infection. In line with this notion, we show that AVH195 modulates autophagy in a balanced manner, without inhibiting the process completely. All observed effects were transient. The effector slows down the development of hypersensitive cell death but does not completely inhibit it. It delays vacuole fusion and autophagic flux but does not fully impair it. Modulations of host autophagy by AVH195 are subtle and do not interfere with vital parameters of the plant cells, so that overexpression of *Avh195* does not generate abnormal phenotypes in plants, and it does not affect the morphology and proliferation potential in Chlamydomonas.

The previously reported involvement of autophagy in ‘pro-survival’ and ‘pro-death’ processes suggests that autophagy plays different, if not opposite roles in responses to biotrophic and necrotrophic pathogens. A study indicated that Arabidopsis mutants defective in different ATG genes display enhanced susceptibility to necrotrophic pathogens, but contrastingly exhibit markedly increased immunity to infection by biotrophic pathogens (Lenz et al., 2011). In the present study, *Avh195* overexpression increases susceptibility to both *P. parasitica* and *H. arabidopsidis*, suggesting that the effector-mediated effect on autophagic flux rather favors biotrophy. The hemibiotrophic strategy of *P. parasitica* has been deciphered during infection of the plant model Arabidopsis (Attard et al., 2010). The biotrophic phase allows the oomycete to successfully establish in the root cortex and to reach the central cylinder, before setting up necrotrophic, invasive growth, and infecting the aerial parts of the plant (Attard et al., 2010). The pathogen then initiates the phase switch between biotrophy and necrotrophy at 15 to 30 h after inoculation. The equilibrium between initial establishment within plant tissues and the development of biomass during invasive growth eventually determines the success of the oomycete life cycle. *Avh195* is expressed during biotrophic growth only, where the effector slows down the development of hypersensitive cell death (a plant response to biotrophs). Once becoming a necrotroph, there is no need any more for the pathogen to interfere with the HR. Taken together, we suggest that AVH195 targets autophagy to decelerate the pro-death function of the mechanism, and thus to promote biotrophy. This interpretation would be in line with the observation that the obligate biotrophic oomycete pathogen *Hpa* develops better in *Avh195*-expressing Arabidopsis than in the Wt.

We show that AVH195 requires the AIMs for slowing down the progression of cryptogein-mediated cell death, as this activity is almost abolished in an effector in which 3 of the 5 AIMs were mutated. We also show that the effector interacts with different ATG8 isoforms from Arabidopsis and the ATG8 from Chlamydomonas. In plant and Chlamydomonas cells expressing *Avh195*, the effector associates with membranes in microsomal fractions, whereas ATG8 occurs in both, soluble and membrane-associated forms. Our co-localization studies support the conclusion that interactions between the effector and ATG8 take place at membrane layers. Functional autophagy depends on the presence of membrane-associated ATG8 proteins, which recruit components of the autophagy machinery as well as cargo to the growing phagophore (Abdollahzadeh et al., 2017). However, ATG8 proteins also modulate biophysical properties of the membranes, and contribute to the fusion of small vesicles for phagophore expansion, and to autophagosome biogenesis (Nakatogawa et al., 2007). Our findings that Chlamydomonas cells accumulate densely charged small vesicles rather than large vacuoles might indicate that the membrane-associated interaction between AVH195 and ATG8 affects either structural aspects of the autophagic machinery, or the administration of potential cargo, or both.

Previously, the *P. infestans* RxLR effector PexRD54 was shown to interact with ATG8 and to target autophagy in its potato host for infection (Dagdas et al., 2016). A PexRD54 gene ortholog is present in the *P. parasitica* genome (PPTG_03663, XP_008894598), but AVH195 and PexRD54 do not have much in common. The two proteins display noticeable differences at the sequence level, only sharing the conserved characteristics of RxLR effectors and the presence of AIMs. Unlike PexRD54, which displays a discriminant, high affinity for potato ATG8Cl, AVH195 does not exhibit isoform-specific binding, despite an apparently higher affinity for AtATG8H than for the two other AtATG8 members that we tested. Furthermore, PexRD54 was proposed to activate autophagy, without disturbing autophagic flux (Dagdas et al., 2016). In the present study, we show that *Avh195* slows down the autophagic process in photosynthetic organisms from 2 distinct kingdoms, instead of activating it. All symptoms we observed fit with the definition of a perturbation of the autophagic flux (Galluzzi et al., 2017). *Avh195* and *PexRD54* thus seem to have distinct specificities and assume distinct functions, while targeting the same mechanism in host cells. As PexRD54 and Avh195 orthologs exist in *P. parasitica*, and *P. infestans*, it would be interesting to analyze how both proteins coordinate interference with host autophagy.

## Materials and methods

### Plant material and oomycete cultures

Arabidopsis lines were from the Colombia (Col-0) genetic background. Plants were grown in growth chambers at 21°C as described (Le Berre et al., 2017). Tobacco (*Nicotiana tabacum* var *xanthii* and *Nicotiana benthamiana*) plants were grown as described (Evangelisti et al., 2013). The *Phytophthora parasitica* strain PPINRA-310 was maintained in the *Phytophthora* collection at INRA, Sophia Antipolis, France. The conditions for *in vitro* growth and zoospore production were as previously described (Galiana et al., 2005). *Hyaloperonospora arabidopsidis* (*Hpa*, isolate Noco2) culture and sporulation were described elsewhere (Hok et al., 2011).

### *Chlamydomonas reinhardtii* culture and transformation

The *Chlamydomonas reinhardtii* strain *dw15.1* (nit1-305 cw15; mt+), a cell wall-less derivative of strain *cw15*, was used for all studies. Seed cells were cultivated in 250-ml flasks with 100 ml Tris Acetate Phosphate (TAP) medium (Harris, 1989) with a 12 h photoperiod at 25°C. Rapamycin (1 mg/ml stock in 90% ethanol-10% Tween 20) was added to cultures at a final concentration of 0.5 µM. Considering the biased, high GC content of the *C. reinhardtii* genome, we designed a synthetic gene encoding *Avh195* using the average codon usage implemented in the Codon Usage Database (http://www.kazusa.or.jp/codon). The gene was synthesized by GeneArt technologies and cloned into the KpnI and NotI restriction sites of the pChlamy_3 vector (Thermo Fisher Scientific). Nuclear transformation was performed using the electroporation method as previously described (Kong et al., 2017a). Briefly, the cells of *Chlamydomonas reinhardtii* strain *dw15.1* were grown to 1.0–2.0 × 10^6^ cells/ml in TAP medium. Subsequently, 2.5 × 10^7^ cells were harvested by centrifugation and suspended in 250 µL of TAP medium supplemented with 40 mM sucrose. Electroporation was performed by applying an electric pulse of 0.7 kV at a capacitance of 50 µF (GENE PULSER, Bio-Rad), using 400 ng of *Sca*<u>I</u>-linearized plasmids purified by agarose gel electrophoresis. Transgenic strains were selected directly on TAP/agar plates containing hygromycin B (30 mg/L), and the plates were incubated under continuous fluorescent light (20 µmol m^−2^ s^−1^) at 25°C before being transferred to the above-described culture conditions.

### Vector construction

Primers used for cloning are listed in Supplementary Table 1. *Avh195* without the signal peptide-encoding sequence was amplified from a full length cDNA clone (Le Berre et al., 2008) with Gateway-compatible primers. Site-directed mutagenesis of *Avh195* was performed using the QuickChange II kit (Stratagene) according to the manufacturer’s recommendations. Three primer pairs were specifically designed to alter selected AIMs (Supplementary Table 1), replacing hydrophobic residues at each site by an alanine. *AtATG8* coding sequences were amplified with Gateway-compatible primers from the plasmids G22544, U17226, and G82070, which were obtained from the Arabidopsis Biological Resource Center (ABRC) at the Ohio State University, and which contain the Arabidopsis genes encoding ATG8D, G, and H, respectively. The *CrATG8* coding sequence was amplified with Gateway-compatible primers from Chlamydomonas *dw15.1*-derived cDNA. For all genes, two sets of amplicons were generated, which contained, or did not contain the Stop codon for C-terminal fusions. All amplicons were recombined in the entry vector pDONR201 (Invitrogen) for further swaps to destination vectors. Destination vectors were pK7FWG2. pK7WGF2 and pK7WGR2 (all from Plant Systems Biology, VIB, Gent) for transient and stable expression in *Nicotiana* and Arabidopsis, respectively, and pMetYC-DEST and pNX32-DEST for Y2H analysis using the mbSUS system (Grefen *et al*., 2009). Entry-to-destination swaps were performed via Gateway LR clonase reactions (Thermo Fisher Scientific), according to the instructions of the supplier.

### Site-directed mutagenesis

Avh195 was examined for the presence of AIMs, otherwise called LIR (LC3-interacting region) motifs (Birgisdottir et al, 2013). This was achieved at the iLIR autophagy database hosted at the https://ilir.warwick.ac.uk. Site-directed mutagenesis of *Avh195* was conducted using a QuickChange II kit (Stratagene) according to the manufacturer’s recommendations. Three primer pairs were specifically designed to alter selected AIMs (see results). This was achieved through the replacement of the hydrophobic residues of each site by an alanine.

### Stable and transient overexpression *in planta*

Arabidopsis plants were transformed using the dipping method (Clough and Bent, 1998) and selected on MS medium plates (1% agar) containing 50 µg/ml kanamycin. Transformed plants were transferred to soil, and seeds were collected. Ten independent primary transformants (T1) harboring the constructs were verified by PCR, and homozygous T3 plants were obtained for further analysis. For transient expression analysis, *A. tumefaciens* strain GV3101 transformed with the various constructs was grown in LB medium supplemented with 50 µg/ml rifampicin, 20 µg/ml gentamicin and 100 µg/ml spectinomycin until OD_600_ reached 1.0. Cells were pelleted, resuspended in infiltration buffer (10 mM MgCl_2_, 10 mM 2-[N-morpholino] ethanesulfonic acid [MES], ph 5.6, 200 µM acetosyringone) and adjusted to an appropriate optical density, then left for 3h at room temperature in the dark before infiltration. The abaxial side of leaves was infiltrated using a syringe without needle. Cell death symptoms were observed over a 72-hour period. For confocal imaging, leaf patches were collected 72 h after infiltration and mounted either in water or in 0.8 M sorbitol prior to analysis.

### Infection assays with oomycetes

For inoculation with *P. parasitica*, *Arabidopsis* seeds were surface-sterilized in 20% NaClO 80% EtOH, rinsed 3 times with 100% ethanol, and sown on MS medium containing 2% sucrose and 2% agar. Seeds were cold-stratified for two days and then maintained at 21°C under short-day conditions (8h light / 16h dark). After 6 days, plantlets were transferred to 96 well plates containing per well 30 µl of the same medium, which was overlaid with 25µl of liquid 0.5x MS containing 1%. Plants were grown for another 8 days under these conditions, before inoculation with 1,000 zoospores per well. All experiments were performed in duplicates, and plantlets that were collected at different time points after inoculation were immediately frozen in liquid nitrogen before analysis by RT-qPCR. Infection assays with *Hpa* isolate Noco2 were performed and sporulation levels were determined as described (Hok et al., 2011). For microscopic analysis of the *Hpa* infection success after 24 h, leaves were collected, stained with Trypan blue according to Hok et al. (2011) and analysed with a Zeiss Axio Zoom.v16 microscope.

### RNA extraction and gene expression analysis

RNA extractions were performed as described (Le Berre et al. 2008) or using the miRNAeasy kit (Qiagen) according to the supplier’s instructions. After DNAse I treatment (Ambion), 1 µg of RNA was reverse-transcribed using the IScript cDNA synthesis kit (BioRad) or Superscript IV RT (Invitrogen). qPCR experiments were performed in an AriaMX (Agilent) thermocycler with 5 *µ*l of a 1:20 cDNA dilution, and 7.5 µl of Brilliant III Ultra-Fast SYBR Green QPCR Master Mix (Agilent), according to the manufacturer’s instructions. Primer pairs for qPCR were designed using Primer3 (http://frodo.wi.mit.edu). Relative expression of the target genes was normalized for Arabidopsis with transcripts from the constitutive genes At5g62050 and At5g11770 (Le Berre et al., 2017) for *P. parasitica* with transcripts from *WS41* and *UBC* (Evangelisti et al., 2013), and for Chlamydomonas with transcripts from Cre02.g106550, Cre04.g227350 and Cre05.g232750 (Zones et al., 2015). Values were either displayed as 2^-ΔΔCt^, or as the normalized relative quantities that were determined by the modified ΔCt method employed by the qBase 1.3.5 software.

### Protein extraction, preparation of microsomes, and immunoblot analysis

Leaf sections from Agrobacterium-infiltrated *Nicotiana* were ground in a mortar under liquid nitrogen. The fine powder was transferred to 15 ml centrifuge tubes and suspended in 100 mM Tris-HCl buffer at pH 7.4 containing 10 mM KCl, 10 mM EDTA, 1% protease inhibitor cocktail, and 12% Sucrose. Samples were centrifuged at low speed (1,500 x *g*) for 10 min at 4°C to pellet tissue debris. Supernatants were collected, filtered through 70 µm cell strainers, and centrifuged at 100,000 x*g* for 90 minutes at 4°C. Supernatants containing soluble proteins were separated from the microsomal pellets, which were washed once with sucrose-free extraction buffer and resuspended in 1 ml of fresh buffer. Microsomal fractions were sonicated on ice at 30% intensity with 3 pulses of 10 seconds each. For protein extraction from *Chlamydomonas* cells, 10 ml of culture treated with either DMSO as the control, rapamycin at 0.5 µM, or Concanamycin A at 0.01 µM for 8 h were collected by centrifugation at 4,000 x *g* for 5min and frozen in liquid nitrogen. Cell pellets were resuspended in 2o0 µl of extraction buffer (50 mM sodium phosphate buffer at pH 5.8 containing 10 mM KCl, 1 mM EDTA, and 1% protease inhibitor cocktail). Cells were lysed with 3 cycles of freeze-thawing between liquid nitrogen and 37 °C, transferred to 1.5 ml reaction tubes, supplemented with Fontainebleau sand and grown with a tube-adapted pistil. The homogenates were centrifuged at 1,000 xg for 8 min at 4 °C, and the recovered supernatants centrifuged in a TLA-120.1 rotor at 120,000 xg for 1 h at 4°C. The supernatants from this centrifugation step were transferred to fresh reaction tubes, and the pellets were washed with 200 ml extraction buffer and dried before 10 µl of 10 % SDS was added to solubilize the microsomes. The samples were then completed with 90 µl extraction buffer. Protein contents in all preparations were determined with the Bradford Ultra dye (Expedeon, Cambridge, UK) according to the manufacturer’s instructions, and samples were adjusted to equal concentrations. Proteins were separated by SDS-PAGE (10 % acrylamide) using the BioRad Mini PROTEAN System, transferred to PVDF membranes (pore size 0.2 µm) with the BioRad Trans-Blot Turbo system, and revealed with antisera directed against ATG8 (Agrisera AS14 2769), GFP (Chromotek 3H9), the photosystem II PsbO proteins (Agrisera AS06 142-33), and FlvB (Chaux et al 2017).

### Yeast two-hybrid analysis

To determine interactions between membrane-associated AVH195 and ATG8, the mating-based split ubiquitin system (mbSUS) was used and employed as described (Grefen et al., 2009). Sequences encoding native or AIM-mutated AVH195 were integrated as baits into pMetYC-DEST and transferred into the haploid Mat_a_ yeast strain THY.AP4. Sequences encoding *Cr*ATG8, *At*ATG8D, *At*ATG8G, and *At*ATG8H, as well as the Arabidopsis receptor IOS1 (Hok et al., 2011) for negative control were cloned as preys into pNX32-DEST and transferred into the haploid Mat_alpha_ yeast strain THY.AP5. This strain was also transformed for positive and negative control with the prey plasmids, pNubWt-X-gate and empty pNX32-DEST, respectively. Mating between THY.AP4 and THY.AP5 transformants, and selection of diploids for growth on Synthetic Complete minimum (SC) medium complemented with adenine (A) and histidine (H) was performed as described (Grefen et al., 2009). Autotrophic growth of yeast cells was determined at 30 °C on SC medium supplemented with 50 µM methionine in the absence of adenine and histidine.

### Live cell imaging

Chlamydomonas cells were stained with the acidotropic dye Lysotracker DND-189 (Thermo Fisher), which was added to cells at 1 µM final concentration. Cells were incubated at 37°C for 30min in the dark, pelleted, rinsed twice with TAP medium, and resuspended in a minimal volume before mounting on glass slides. Cells were analyzed with the epifluorescence microscope AxioimagerZ1 (Zeiss) using the GFP44 filter. Images were acquired with an AxioCam MRm camera and analyzed with the Zeiss ZEN 2102 pro and Fiji (https://imagej.net/Fiji) image processing softwares. Green and red fluorescence conferred by GFP- and RFP-tagged fusion proteins were detected in optical sections by confocal laser scanning microscopy on an inverted Zeiss LSM 880 microscope, equipped with Argon ion and HeNe lasers as excitation sources. For simultaneous imaging of GFP and RFP, samples were excited at 488 nm for GFP and 561 nm for RFP. Confocal images were processed using the Zeiss ZEN 2 software.

### Transmission Electron Microscopy

Cells from Wt and transgenic Chlamydomonas lines were cultured in liquid medium until the exponential growth phase (2 x 10^6^ cells/ml) was reached under a 12h light/12h dark cycle. When appropriate, rapamycin was added to 0.5 µM final concentrations at the beginning of the light period. After 4, 8, 12 and 24 hours of incubation, cells were collected, fixed in a mixture of cacodylate buffer 0.1M and glutaraldehyde 2.5% and stored at 4°C. Cells where rinsed with buffer then postfixed in 1% osmium tetroxide in cacodylate buffer (0.1 M). The final cell pellets were washed in water, dehydrated in acetone, and embedded in epoxy resin. Uranium- and lead citrate-contrasted thin sections (80 nm) were analyzed in a JEOL JEM-1400 120kV transmission electron microscope. Images were acquired with an 11 MegaPixel SIS Morada CCD camera (Olympus).

### Flow cytometric analyses

Analyses by flow cytometry were performed on a SP6800 spectral cytometer (SONY Biotechnologies (Futamura et al., 2015). Chlamydomonas cells were labeled with 7.5 µM CFSE (Life technologies; 2×10e7 cells/ml) for 20 minutes at 37°C, then washed once in TAP medium and resuspended in appropriate amounts of fresh medium to reach final concentrations of 1 x 10^6^ cells/ml. CFSE-labeled cells were then treated with 0.004% EtOH and 0.001% Tween, or with rapamycin at 0.5 µM. Assessing the beginning of the light period as time point 0, aliquots were taken from the cultures 4, 8, 12 and 24h later. For each time point, at least 80.000 cells were collected and analyzed for: i) size and overall complexity (FSC and SSC parameters), ii) chlorophyll content (natural autofluorescence collected after excitation with the 488 and 405 nm laser lines), and iii) proliferation based on CFSE dilution. Data from single cells were then analyzed with the Kaluza software (Beckman coulter). The dynamics of parameter changes were estimated by measuring the values collected at T_N_ reported to the basal values at the beginning of the kinetics (T_0_).

## Accession numbers

*Avh195*: FK938647; PpNPP1: AAK19753; PpHMP1: XP_008904491; *Pp*WS41: CF891677; *Pp*UBC: CK859493; *Cr*ATG8: XP_001699190; *Cre*02.g106550: XM_001699695; *Cre*04.g227350: XM_001692180; *Cre*05.g232750: XM_001702168; CrATG8: XP_001699190; AtATG8D: NP_178631; AtATG8G: NP_191623; AtATG8H: NP_566283.

## Acknowledgements

We would like to thank Claire Venault-Fourrey (INRA, Nancy, France), Eric Galiana (INRA, Sophia Antipolis, France), Elodie Gaulin and Christophe Roux (Toulouse University-CNRS, France) for fruitful discussions. Live cell imaging work was performed at the SPIBOC imaging facility of the Institut Sophia Agrobiotech. We thank Dr Olivier Pierre and the whole platform team for their help with microscopy. This work was supported by the French Government (National Research Agency, ANR) through the “Investments for the Future” LABEX SIGNALIFE: program reference # ANR-11-LABX-0028-0, Plant-KBBE project dsRNAguard, the A*MIDEX (ANR-11-IDEX-0001-02) and the ANR JCJC MUsCA (ANR-13-JSV5-0005) projects. The authors also acknowledge the European Union Regional Developing Fund, the Region Provence Alpes Cote d’Azur, the French Ministry of Research, and the CEA for funding the HelioBiotec platform at CEA Cadarache. The authors declare no potential conflicts of interest.

## Authors contributions

FP, HK and MLK conceived the study. ST, MLK and VA carried out experiments on Chlamydomonas transformants and plants. PA, FK and GP generated Chlamydomonas transformants. SP and JC managed electron microscopy and flow cytometry analyses, respectively. ST, MLK, JC, HK and FP analyzed the data. HK and FP wrote the manuscript. All authors read and approved the final manuscript.

## Online Supplemental material

Fig. S1 shows flow cytometry analyses of Chlamydomonas transformant cell lines in normal conditions. Fig. S2 shows flow cytometry analyses of the same transformant lines upon rapamycin treatment. Fig. S3 shows TEM images for accumulation of lysosome-like structures in Avh195-overexpressing Chlamydomonas lines. Fig. S4 shows typical TEM images for Chlamydomonas cultures untreated or treated with rapamycin. Fig. S5 shows a typical infection of Wt and transgenic Arabidopsis leaves by *Hyaloperonospora arabidopsidis*, indicating that the pathogen develops faster in lines expressing *Avh195*. Table S1 shows a list of primers used in this study.

## Supplementary data

**Supplementary Figure S1: Flow cytometry analyses of Chlamydomonas transformant cell lines.**

*Chlamydomonas* cultures of Wt and transformant strains were analyzed over a period of 24 hours. For each time point, at least 80.000 cells were controlled for the following parameters: Forward light scattering - FSC (A); Side light scattering - SSC (B); Autofluorescence of cells (C), indicative for chlorophyll content; CFSE repartition into daughter cells (D), here presented as CFSE^-1^ to highlight cell proliferation. Plots on the left represent median values of fluorescence intensity (arbitrary units) over a 24 h period. Plots on the right represent the distribution of fluorescence intensity within the indicated population of cells collected at time point 8 h.

**Supplementary Figure S2: Flow cytometry analysis of Chlamydomonas transformant cell lines upon rapamycin treatment.**

*Chlamydomonas* cultures of Wt and transformant strains were analyzed as described in Supplementary Figure S1.

**Supplementary Figure S3: Accumulation of electron-dense, lysosome-like structures in cells from *Avh195*-expressing Chlamydomonas lines that were not treated with rapamycin.**

High resolution TEM micrographs were recorded at different time points after onset of light over a 24 h day/night cycle. Appearance of vesicles likely reflects basal autophagic flux within the cells. Bars represent 1 µm.

**Supplementary Figure S4: Representative view of Chlamydomonas cells from the wild-type and transgenic lines expressing *Avh195*, as analysed by TEM.**

Micrographs show untreated cells, or cells that were incubated with 0.5 µM rapamycin for 4 h, 8 h, 12 h, and 24 h. Bars represent 10 µm.

**Supplementary Figure S5: Micrograph overview of *H. arabidopsidis*-infected leaves.**

Leaves from the Wt and from Avh195-expressing Arabidopsis lines OE6 and OE9 were collected 24 h after inoculation with *Hpa*, stained with trypan blue and analyzed with an Axio Zoom.v16 microscope. Bars represent 100 µM.

**Supplementary Table 1.** Primers used in this study.

