## Supplemental data for "An oomycete effector impairs autophagy in evolutionary distant organisms and favors host infection"

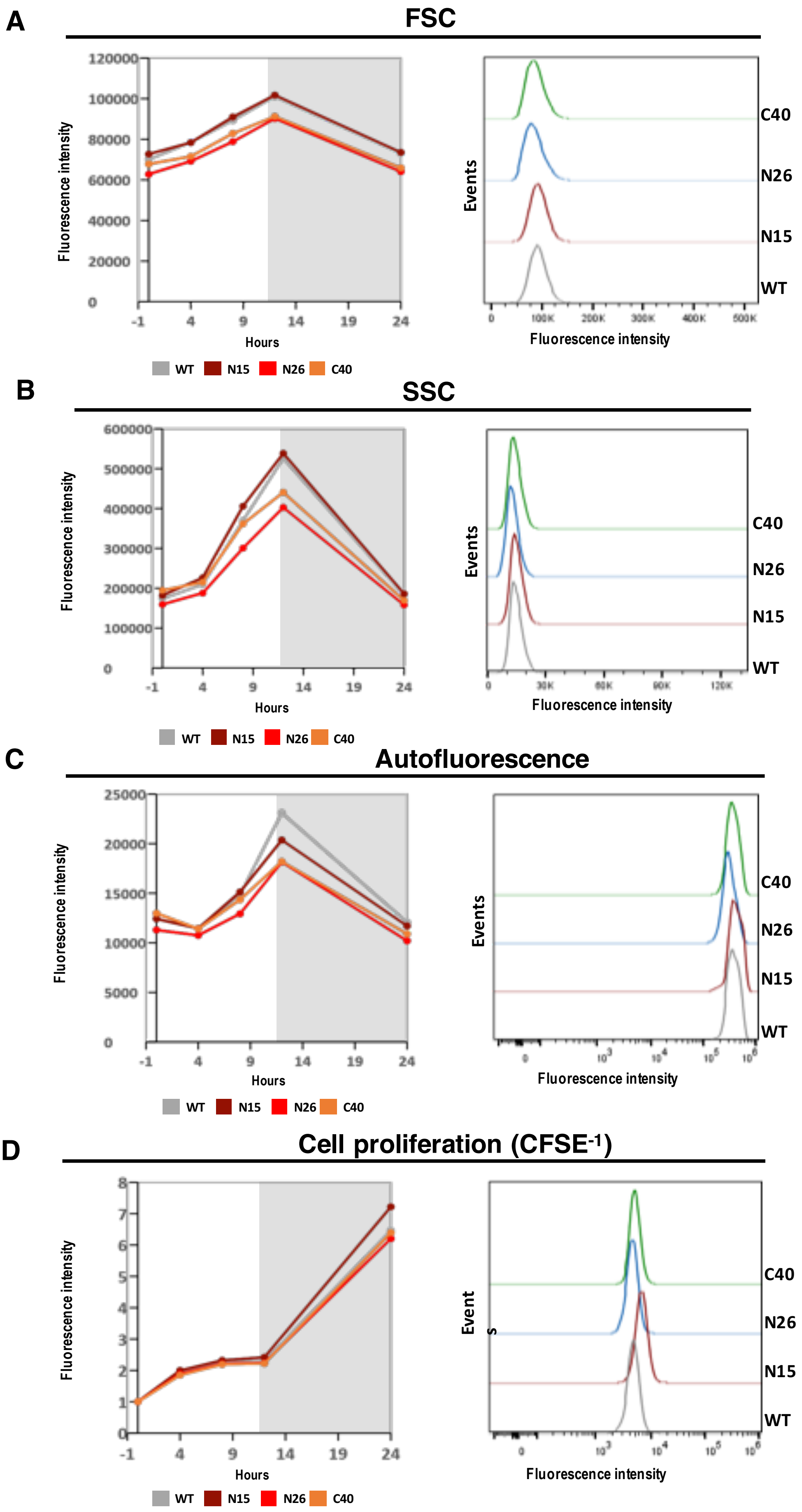
A
FSC
B
SSC
C
Autofluorescence
D
Cell proliferation (CFSE-1)
Fluorescence intensity
Hours
Events
Fluorescence intensity
C40
N26
N15
WT
WT N15 N26 C40

**Supplementary Figure S1: Flow cytometry analyses of Chlamydomonas transformant cell lines.**
*Chlamydomonas* cultures of Wt and transformant strains were analyzed over a period of 24 hours. For each time point, at least 80.000 cells were controlled for the following parameters: Forward light scattering - FSC (A); Side light scattering - SSC (B); Autofluorescence of cells (C), indicative for chlorophyll content; CFSE repartition into daughter cells (D), here presented as $CFSE^{-1}$ to highlight cell proliferation. Plots on the left represent median values of fluorescence intensity (arbitrary units) over a 24 h period. Plots on the right represent the distribution of fluorescence intensity within the indicated population of cells collected at time point 8 h.

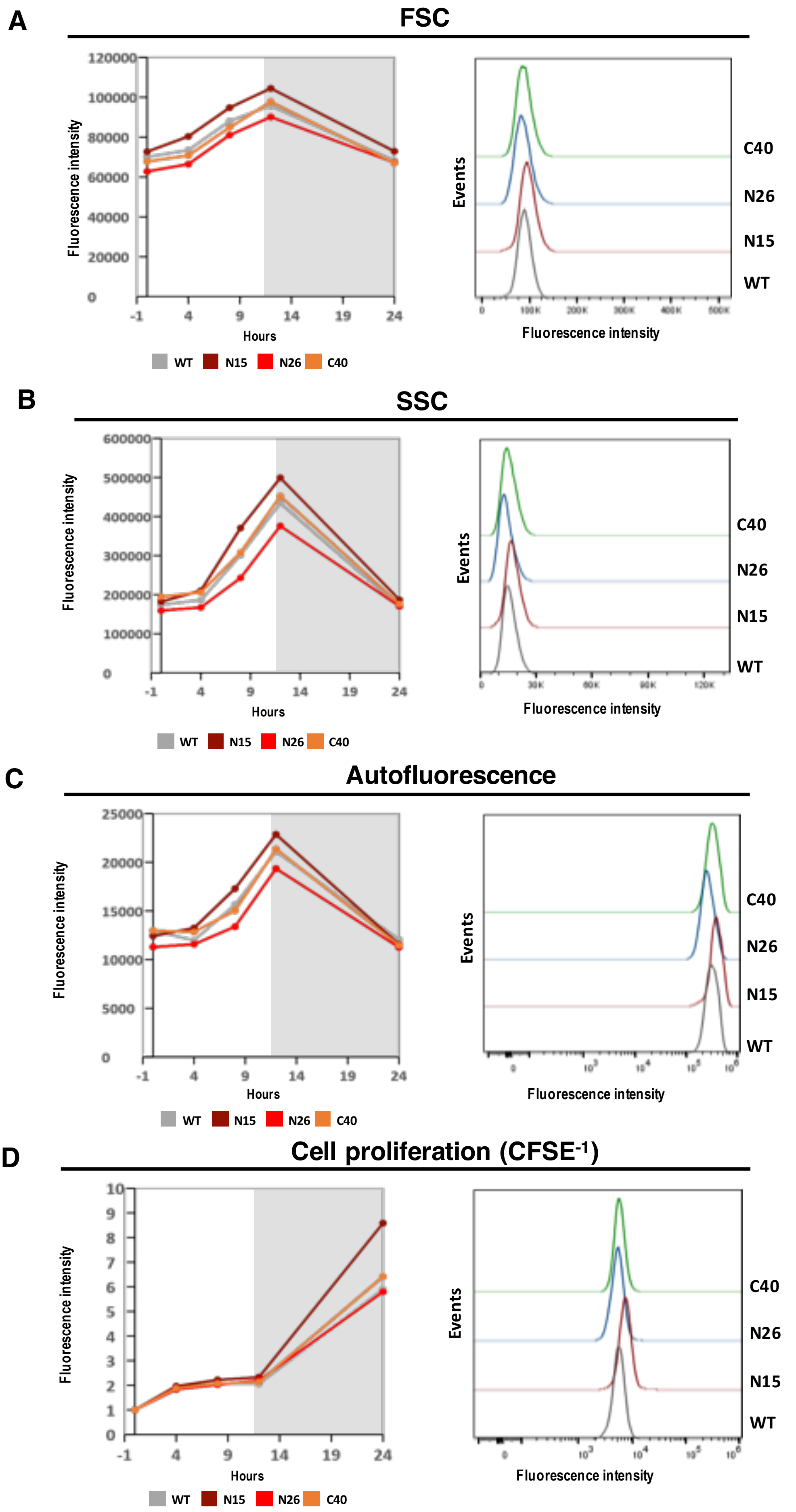


**Supplementary Figure S2: Flow cytometry analysis of Chlamydomonas transformant cell lines upon rapamycin treatment.** *Chlamydomonas* cultures of Wt and transformant strains were analyzed as described in Supplementary Figure S1.

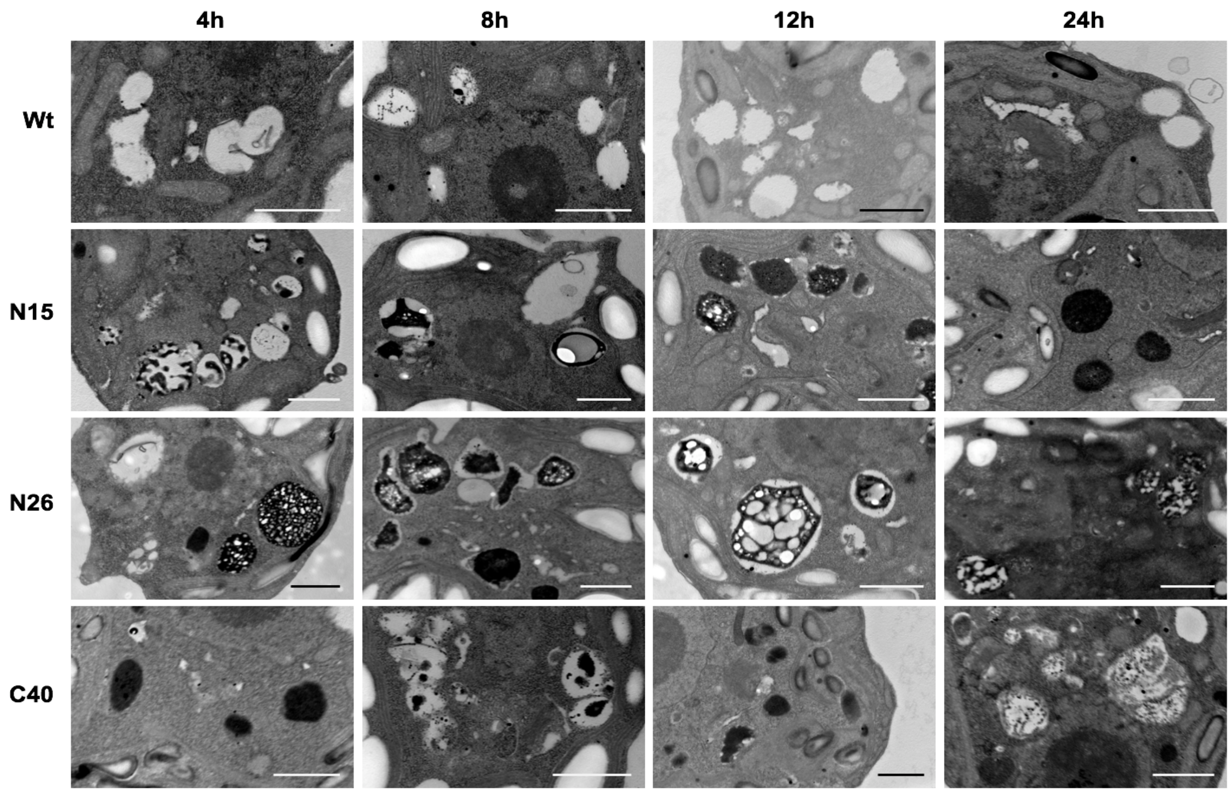


**Supplementary Figure S3: Accumulation of electron-dense, lysosome-like structures in cells from *Avh195*-expressing Chlamydomonas lines that were not treated with rapamycin.**

High resolution TEM micrographs were recorded at different time points after onset of light over a 24 h day/night cycle. Appearance of vesicles likely reflects basal autophagic flux within the cells. Bars represent 1 µm.

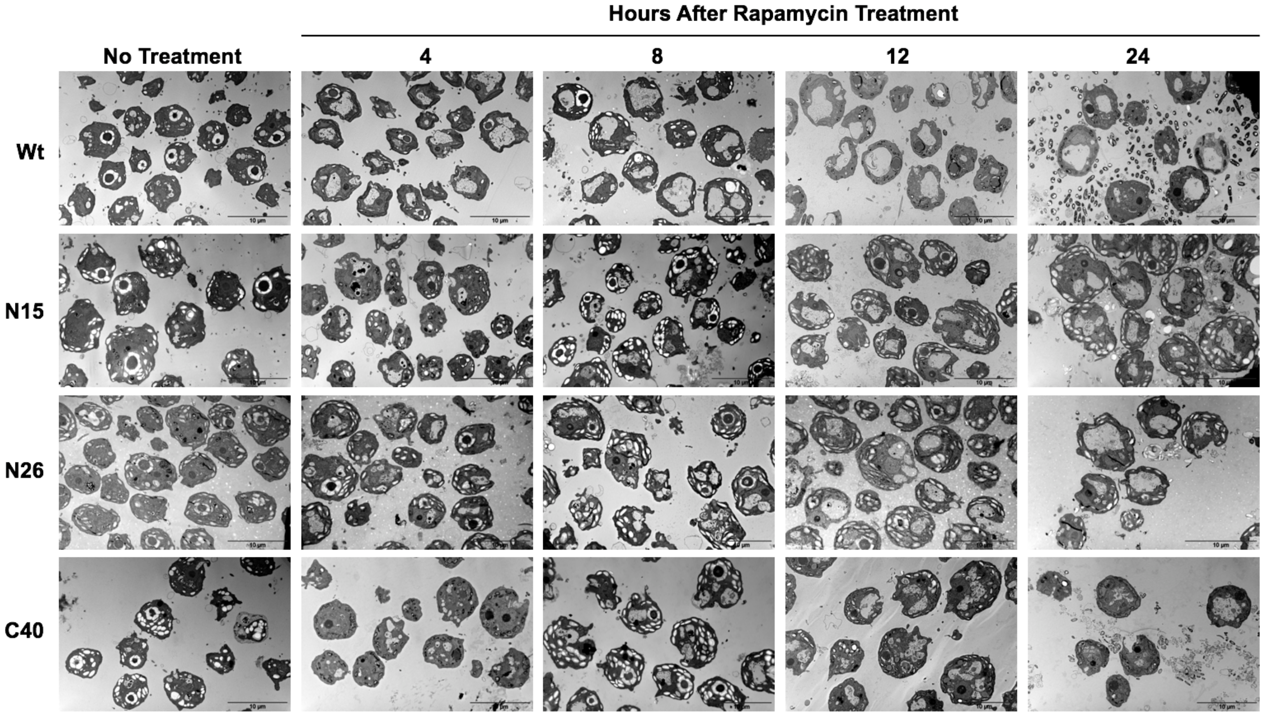


**Supplementary Figure S4: Representative view of Chlamydomonas cells from the wild-type and transgenic lines expressing *Avh195*, as analysed by TEM.**

Micrographs show untreated cells, or cells that were incubated with 0.5 µM rapamycin for 4 h, 8 h, 12 h, and 24 h. Bars represent 10 µm.

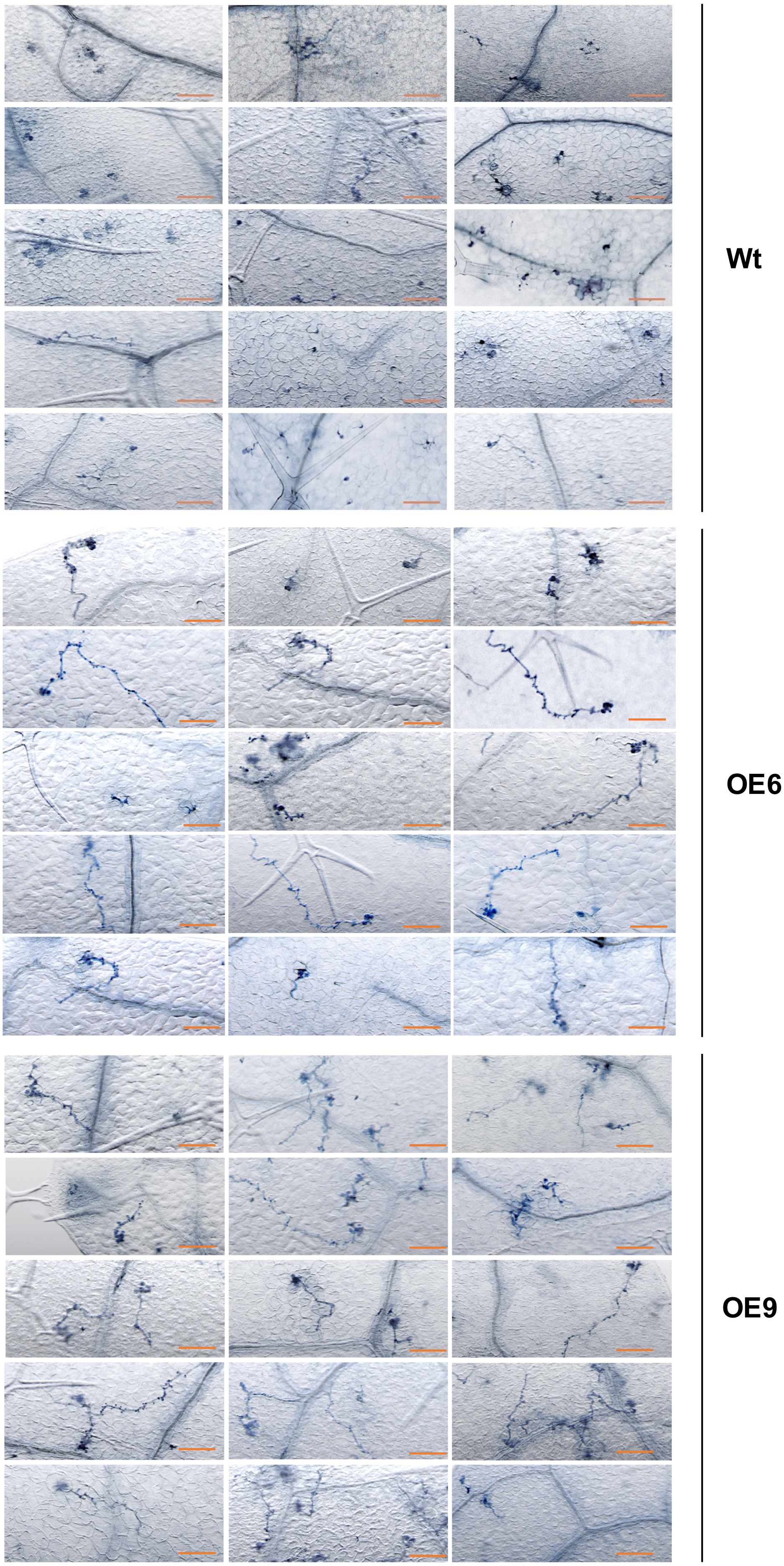


**Supplementary Figure S5: Overview screen of *Hpa* infection sites on Arabidopsis leaves.**
Micrographs of 15 trypan blue-stained infection sites on leaves from each the Wt and the transgenic *Avh195*-expressing lines OE6 and OE9, 24 h after inoculation. Developing hyphae grow overall faster in tissue of the transgenic lines. Bars represent 100 µm.

**Supplementary Table 1**: Primers used in this study.

| **Name** | **Organism** | **Sequence 5'3'** | **Comment** |
|---|---|---|---|
| attB1_Avh195_F | *P. parasitica* | GGGGACAAGTTTGTACAAAAAAGCAGGC TTCACCATGCTATCGGCCTATGAACAA | Gateway cloning |
| attB2_Avh195_R | *P. parasitica* | GGGGACCACTTTGTACAAGAAAGCTGGG TCTCACAAGGCGCTAGCCGCGTG | Gateway cloning |
| attB2_Avh195_ΔEER_R | *P. parasitica* | GGGGACCACTTTGTACAAGAAAGCTGGG TCTCACCTCTCTTCAGCATC | Gateway cloning |
| 195_AIM_S1_mut_F | *P. parasitica* | GAAAAACGGGAAAAGCGCTGATGACGCC TTCGACCGCTGG | Site_directed mutagenesis |
| 195_AIM_S1_mut_R | *P. parasitica* | CCAGCGGTCGAAGGCGTCATCAGCGCTTT TCCCGTTTTTC | Site_directed mutagenesis |
| 195_AIM_S2_mut_F | *P. parasitica* | CATCTTCGACCGCGCGATTCGAGCCGATA AGTCACCG | Site_directed mutagenesis |
| 195_AIM_S2_mut_R | *P. parasitica* | CGGTGACTTATCGGCTCGAATCGCGCGGT CGAAGATG | Site_directed mutagenesis |
| 195_AIM_S3_mut_F | *P. parasitica* | CAATCAGACCGATTGCGCGCGAAGCCGG ACTGACAGAG | Site_directed mutagenesis |
| 195_AIM_S3_mut_R | *P. parasitica* | CTCTGTCAGTCCGGCTTCGCGCGCAATCG GTCTGATTG | Site_directed mutagenesis |
| attB1_CrATG8_F | *C. reinhardtii* | GGGGACAAGTTTGTACAAAAAAGCAGGC TTCACCATGGTTGGCTCCCGACCCCCGAC | Gateway cloning |
| attB2_CrATG8_R | *C. reinhardtii* | GGGGACCACTTTGTACAAGAAAGCTGGG TCTCACAACGCCAGCTCCTCCACA | Gateway cloning |
| pChlamy3_intron_F | *C. reinhardtii* | TGCTTGCAGATTTGACTTGC | Chlamyodomans genoytping |
| pChlamy3_spliced_F | *C. reinhardtii* | TAAAATGGCCAGGAGATTCG | Chlamyodomans genoytping |
| pChlamy3_RBCS2 3'UTR_R | *C. reinhardtii* | TACCGCTTCAGCACTTGAGA | Chlamyodomans genoytping |
| pChlamy3_5'UTR_F | *C. reinhardtii* | GATAAACCGGCCAGGGGGCC | Chlamyodomans genoytping |
| pChlamy3_3'UTR_R | *C. reinhardtii* | CAGCAAAAGGTAGGGCGGGC | Chlamyodomans genoytping |
| AT5G11770 _F | *A.thaliana* | GAAGTTGTGCCAATGGAGGT | qPCR |
| AT5G11770 _R | *A.thaliana* | CCACCAATGCAAGAAATCCT | qPCR |
| AT5G62050 _F | *A.thaliana* | AACAGGACTCAGCGATGTTG | qPCR |
| AT5G62050_ R | *A.thaliana* | TACCTGATCTGCCTCCACCT | qPCR |
| qUBC_F1 | *P. parasitica* | CCACTTAGAGCACGCTAGGA | qPCR |
| qUBC_R1 | *P. parasitica* | TACCGACTGTCCTTCGTTCA | qPCR |
| qWS21_F1 | *P. parasitica* | CTCCAGAACGTGTACATCCG | qPCR |
| qWS21_R1 | *P. parasitica* | TAGCGCCCTTCTCCTCAG | qPCR |
| qWS41_F1 | *P. parasitica* | TTCAAGTCCAGTGAGATCGG | qPCR |
| qWS41_R1 | *P. parasitica* | TTGTGTCTTTGTGTGATGCG | qPCR |
| q195_F10 | *P. parasitica* | AGGCAAGCAGCCAAAAAC | qPCR |
| q195_R10 | *P. parasitica* | CGGCACGAAGTTGATACTCTG | qPCR |
| q195_F2 | *P. parasitica* | CTTCGTGCAATGCTCTATCG | qPCR |
| q195_R2 | *P. parasitica* | CAGACGTATCTCCGGTTTCAG | qPCR |
| qNPP1_F | *P. parasitica* | CCCCAAATGAACGTCCTTAC | qPCR |
| qNPP1_R | *P. parasitica* | TGAACTTGACACCAGCCTTC | qPCR |
| qHMP1_F | *P. parasitica* | GATCGGTGAGACCATTTTCG | qPCR |
| qHMP1_R | *P. parasitica* | TGTTGAGGAACGTGTCAAGC | qPCR |
| qCre195_1_F | *C. reinhardtii* | ATCTTCGACCGCTGGATTC | qPCR |
| qCre195_1_R | *C. reinhardtii* | TGGTCTCCAGGTTCATGTTG | qPCR |
| CBLP_F | *C. reinhardtii* | GCCACACCGAGTGGGTGTCGTGCG | qPCR |
| CBLP_R | *C. reinhardtii* | CCTTGCCGCCCGAGGCGCACAGCG | qPCR |
| RBCS2_F | *C. reinhardtii* | ATACTGCTCTCAAGTGCTGAAGCG | qPCR |
| RBCS2_F | *C. reinhardtii* | AAAGACTGATCAGCACGAAACGG | qPCR |
